# From Generalist to Specialist: Evolution of PS2 α-integrins and Implications for Drug Targeting

**DOI:** 10.64898/2026.04.29.721650

**Authors:** Shixi Sun, Yunfeng Chen, Rong-Guang Xu, Heng Zhang, Fahad Mostafa, Li Liu

## Abstract

Integrins are heterodimeric transmembrane receptors that mediate cell–cell and cell– extracellular matrix interactions and play essential roles in development and disease. Within the PS2 α-integrin subfamily, four paralogs (αIIb, α5, α8, and αV) share a conserved RGD-binding motif yet exhibit diverse functional specializations. Integrins have been widely targeted therapeutically for various clinical conditions, though achieving subtype specificity remains a major challenge. Here, we performed an integrative evolutionary analysis of 114 PS2 α-integrin sequences across 28 vertebrate species, combining phylogenetic reconstruction, time calibration, ancestral sequence inference, and structural mapping. Our time-calibrated phylogeny indicates that the PS2 lineage originated ∼862 Mya, with diversification of the four paralogs occurring prior to vertebrate radiation. Ancestral state reconstruction reveals that fibronectin and vitronectin binding are ancestral traits, whereas fibrinogen binding and β3 pairing arose independently in the αIIb and αV lineage. Evolutionary rate analysis shows domain-specific divergence, with the β-propeller acting as a hotspot of evolutionary change, likely driven by combined pressures from ligand binding and β-subunit interaction. These pressures vary across paralogs: αIIb exhibits accelerated evolution in ligand-binding regions, while αV displays elevated rates in β-subunit interaction domains. Mapping sequence variation onto structural interfaces identifies lineage-specific substitutions underlying functional divergence, including distinct molecular solutions for fibrinogen binding in αIIb and αV. These findings collectively demonstrate that PS2 α-integrins evolved from a generalist ancestor through neofunctionalization and lineage-specific specialization. This work provides an evolutionary framework for identifying subtype-specific functional sites and highlights the potential of evolution-informed strategies to guide the development of more selective integrin-targeting therapeutics.

## Introduction

Integrins are cell surface receptors that coordinate mechanical and chemical signaling between a cell and its microenvironment [1, 2]. The human integrin family comprises 24 heterodimers formed through noncovalent pairing of one of 18 α-subunits with one of 8 β-subunits. Both the α- and β-subunits are type-I transmembrane proteins composed of a N-terminus ectodomain that binds components of the extracellular matrix (ECM) or counter-receptors on other cells, a single-pass transmembrane helix, and a C-terminus cytoplasmic domain that interacts with cytoskeletal and signaling proteins [3]. These heterodimers exhibit distinct ligand-binding specificities and cell-type distributions, enabling diverse and context-dependent biological functions [4]. Aberrant integrin expression or signaling contributes to a broad spectrum of human diseases, including thrombosis, cardiovascular disease, cancer, immune dysregulation, and inflammatory and fibrotic disorders [5]. Consequently, integrins have emerged as important therapeutic targets [6, 7].

In this study, we focus on the PS2 α-integrin subfamily, comprising four members—αIIb, α5, αV, and α8 [8]. When paired with their corresponding β-subunits, all members in this subfamily recognize the Arg-Gly-Asp (RGD) motif found in ECM proteins such as fibronectin (Fn), vitronectin (Vn), and fibrinogen (Fb). Despite this shared ligand-recognition mechanism, PS2 α-integrins exhibit striking divergence in tissue-specific functions [2]. For instance, αIIb is a platelet/thrombocyte-restricted receptor that mediates Fb-dependent clot formation; α5 is the primary Fn receptor involved in matrix assembly; αV is a broadly expressed integrin central to angiogenesis and tumor biology, contributing to tissue repair, TGF-β activation, tumor progression, and fibrotic disease; and α8 is a tissue-restricted receptor important in developmental and organ-specific adhesion, including roles in fibroblast differentiation. Drugs targeting this family were among the first successful integrin-directed therapies, including αIIbβ3 antagonists such as abciximab, eptifibatide, and tirofiban for thrombosis [9], as well as αV-targeting agents such as cilengitide for cancer [10]. However, significant challenges remain, including excessive bleeding, limited subtype selectivity, compensatory signaling among paralogs, and context-dependent therapeutic responses [6, 7]..

Evolutionary analysis of integrin sequences across diverse species provides a powerful framework to understand structural conservation, functional divergence, and lineage-specific adaptations. While previous studies have established the fundamental phylogenetic relationships of integrins, these analyses were conducted prior to the availability of large-scale genomic datasets spanning diverse taxa, thus having limited species coverage and incomplete representation of paralogs across species [1, 11, 12]. Furthermore, few investigations have integrated these evolutionary signals with the functional constraints governing ligand-binding specificity and cell-type–specific signaling.

In this study, we assembled a comprehensive set of PS2 α-integrin sequences from 28 vertebrate species, spanning over 450 million years of history from cartilaginous fish to primates [13]. We not only reconstructed sequence-level phylogeny but also inferred ancestral functional traits across the evolutionary trajectory. By integrating phylogenetic reconstruction, ancestral sequence inference, and structural mapping, we delineated patterns of conservation and divergence within the PS2 α-integrin subfamily. Our results further reveal a transition from a broadly functional, generalist ancestral integrin to lineage-specific specialist paralogs with distinct structural and functional adaptations. By linking evolutionary constraints to functional architectures, particularly ligand-binding interfaces and β-subunit pairing, we identified sequence variations associated with neofunctionalization and lineage-specific specialization. This evolution-informed approach is broadly applicable to other complex protein families and establishes a framework for rational target selection and the development of more selective and effective therapeutics.

## Materials and Methods

### Sequence collection

Protein sequences of PS2 α-integrins were retrieved from the National Center for Biotechnology Information (NCBI) database [14] (last accessed in April 2026). To ensure comparability across paralogs, a consistent set of 28 vertebrate species was selected for each paralog (αIIb, α5, αV, and α8), spanning diverse taxonomic groups including mammals, birds, amphibians, teleost fish, and cartilaginous fishes (**Fig. 1A**). Sequences from *D. melanogaster* (fruit fly) and *C. elegans* (roundworm) were included as outgroups to root the phylogenetic analyses. Because signal peptide, transmembrane, and cytoplasmic regions are known to align poorly [11], these segments were excluded, and only the mature ectodomain region was retained. In total, 114 sequences (28 × 4 plus 2 outgroups, **Supplementary Table 1**) were used for subsequent phylogenetic and functional analyses (**Fig. 2B**).

**Figure 1.**
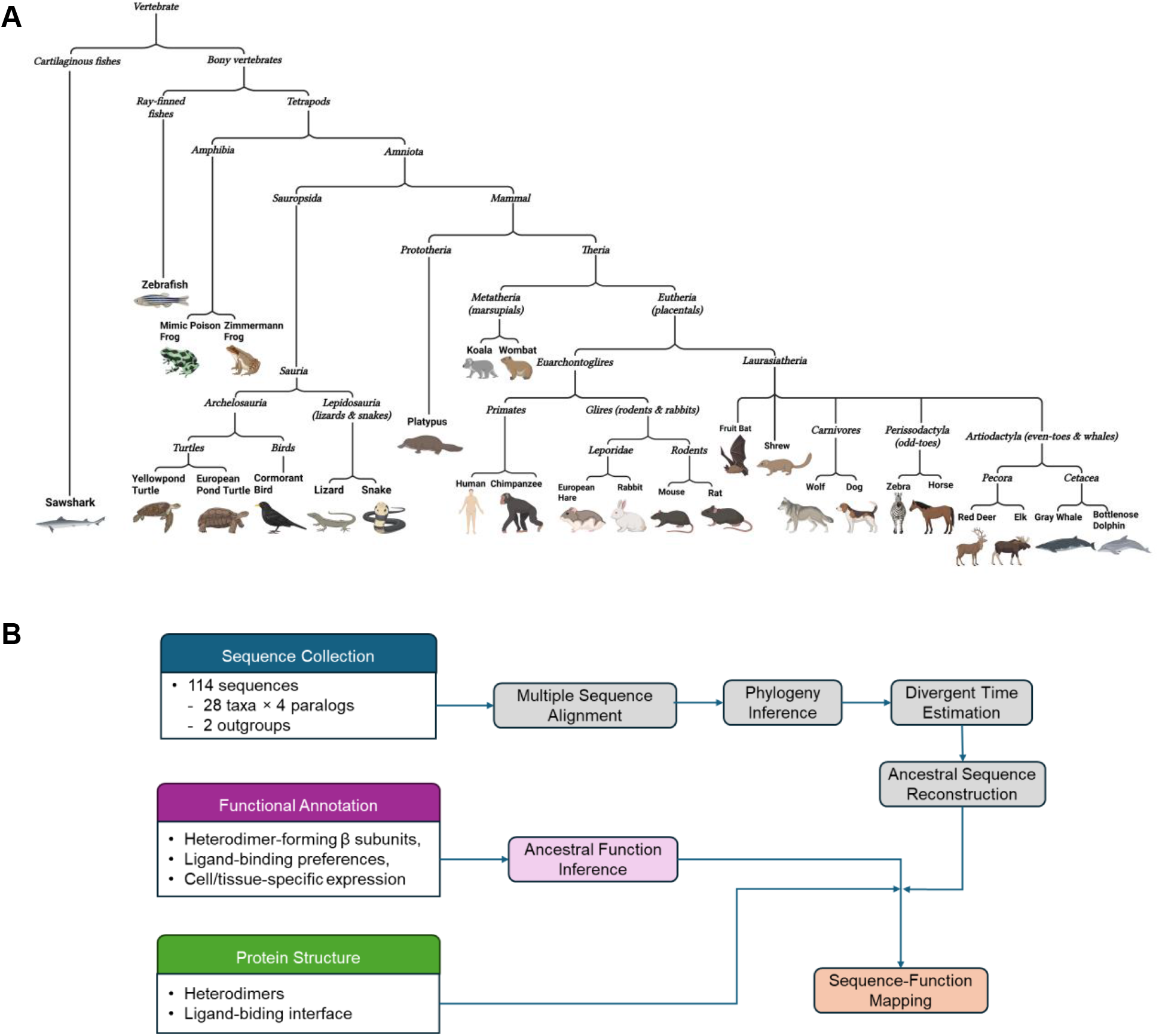
Overview of species selection and analysis workflow. (**A**) Taxonomic distribution of the 28 vertebrate species used in this study. (**B**) Analytical pipeline of the phylogenetic and functional analyses.

**Figure 2.**
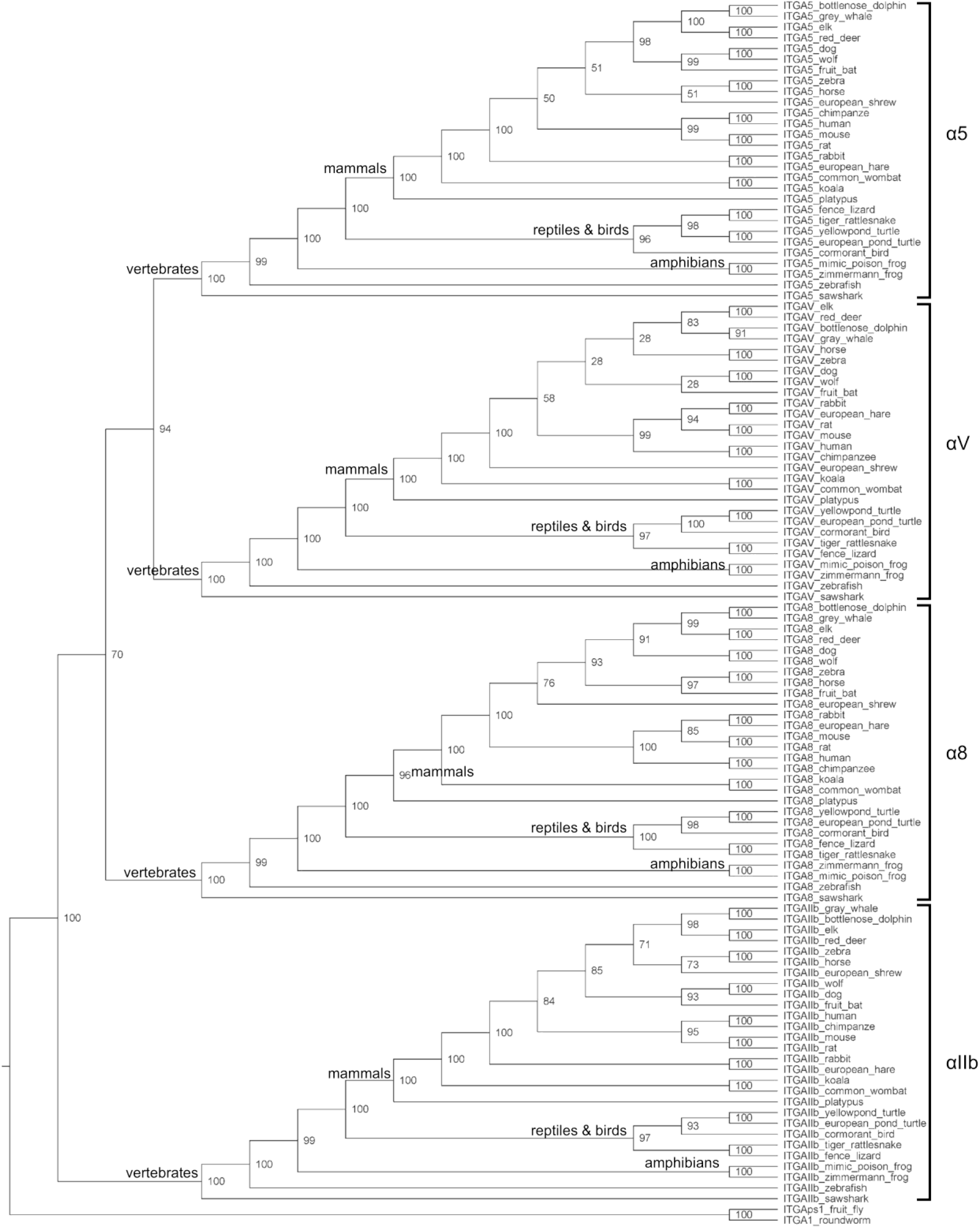
Phylogenetic tree of PS2 α-integrins. Branch labels indicate percentage of 1000 bootstrap pseudo-samples supporting that branch. Major clades corresponding to each integrin paralog are indicated on the right. Major taxonomic groupings are annotated within each clade.

### Phylogenetic analysis

Multiple sequence alignment (MSA) was performed using MAFFT to detect optimal global homology [15] (parameters: BLOSUM62 scoring matrix, G-INS-1 in the progressive step, G-INS-i in the iterative refinement step, default guide tree). The alignment was visually inspected to confirm consistency across conserved regions and to identify potential misalignments or gap artifacts. Based on the MSA, the phylogenetic tree topology was constructed using the Maximum Likelihood (ML) method implemented in MEGA [16] (parameters: 1000 bootstraps, JTT model, 5-category discrete Gamma distribution). Divergence times were estimated using a Bayesian Markov Chain Monte Carlo (MCMC) framework implemented in BEAST [17] (parameters: 10 million MCMC generations, JTT model, relaxed log-normal molecular clock). Fossil calibration was applied at the root (627 Mya based on divergence between human and fruitfly as recorded in TimeTree [18]. Ancestral sequences (Anc) for all internal nodes were reconstructed using the ML framework implemented in PAML [19] (parameters: JTT model, Gamma distribution). In the postprocessing of PAML results, for positions containing only gaps within a given clade, the corresponding ancestral state was assigned as a gap. For positions where the highest posterior probability was below 0.6, ancestral residues were retained but reported in lowercase to indicate low confidence. Given the MSA and the time-calibrated phylogenetic tree, site-specific evolutionary rates were estimated using MEGA (5-category discrete Gamma distribution) [16].

### Inference of ancestral functional states

Based on literature reviews [6, 8, 11], human PS2 α-integrins were annotated with binary functional states on heterodimer formation, ligand-binding preference, and tissue expression patterns. These annotations were used as proxies for each paralog across extant vertebrates. Ancestral functional states were then inferred along the phylogenetic tree based on maximum parsimony (MP).

## RESULTS

### PS2 α-integrin emerged early in eukaryote evolution and diversified among vertebrates

With *D. melanogaster* and *C. elegans* serving as outgroups, the ML phylogenetic tree of PS2 α-integrins shows that vertebrate PS2 α-integrins form a distinct monophyletic group separated from invertebrate homologs, indicating that the core PS2 lineage was established before the emergence of vertebrates. Within vertebrates, the PS2 subfamily diversified into four well-defined clades corresponding to αIIb, α5, αV, and α8 paralogs across 28 species (**Fig. 2**). Each paralog forms a monophyletic cluster with strong bootstrap support (100%), indicating clear evolutionary separation. Among these, α5 and αV group more closely together, suggesting a more recent common ancestry relative to α8, while αIIb forms the most distinct and basal clade. Within each paralog, species relationships generally follow known vertebrate taxonomy, with mammals, reptiles, amphibians, teleost fish, and cartilaginous fish clustering as expected. Tthese results support a stepwise divergence model of PS2 integrins and highlight both conserved lineage structure and paralog-specific evolutionary trajectories.

Time calibration of the phylogenetic tree using a Bayesian MCMC framework running 10 million generations indicates that the PS2 α-integrin lineage emerged approximately 862 million years ago (Mya), a period when eukaryotes were becoming established. This is consistent with the presence of integrin-like proteins in basal eukaryotes such as algae [20]. The most recent common ancestor (MRCA) of vertebrate PS2 α-integrins is estimated at ∼762 Mya, within the broader time frame associated with the early evolution of multicellular animals. Diversification of the four paralogs was completed prior to early vertebrate diversification (∼500 Mya). Because orthologs within each paralog clade exhibit consistent phylogenetic relationships that recapitulate known taxonomy, we collapsed each clade to generate a reduced phylogeny of paralogs (**Fig. 3**). Using the same set of 28 vertebrate species makes the estimated divergence times comparable across the four paralogs. The Anc of α5, αV, and α8 predates that of αIIb. However, the αIIb lineage solidified its identity the earliest at approximately 568 Mya, while αV is the most recently diverged paralog at approximately 494 Mya. Independent analyses using 50 million MCMC generations produced highly consistent divergence time estimates, supporting the robustness of these results.

**Figure 3.**
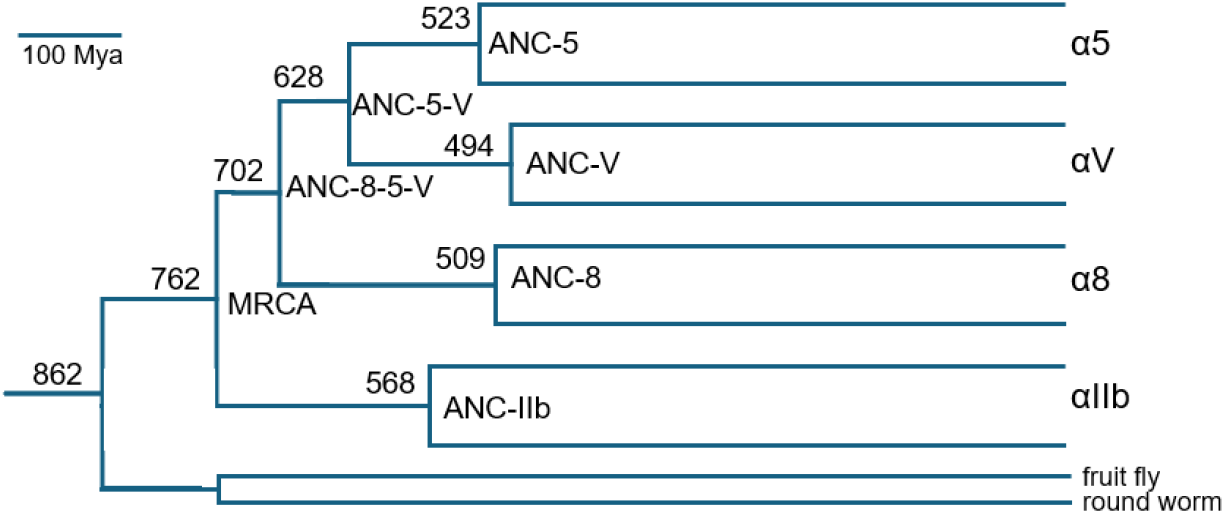
Time-calibrated phylogeny of PS2 α-integrin paralogs. Orthologs within each paralog clade are collapsed. Node labels indicate divergence times (Mya), and annotated nodes represent ancestral duplication events.

### Heterogeneous evolutionary rates reveal region- and paralog-specific divergence

The α-integrin ectodomain consists of a globular headpiece supported by an elongated stalk (**Fig. 4A**). The headpiece adopts a characteristic seven-bladed β-propeller structure, with each blade composing approximately 60 amino acids (repeat R1 – R7, **Fig. 4B**). The stalk region includes the thigh domain and two calf domains (calf-1 and calf-2), which together provide structural support and enable conformational changes associated with integrin activation [6]. To investigate patterns of conservation and divergence across PS2 α-integrins, we estimated site-specific evolutionary rates and mapped these rates onto the structural domains.

**Figure 4.**
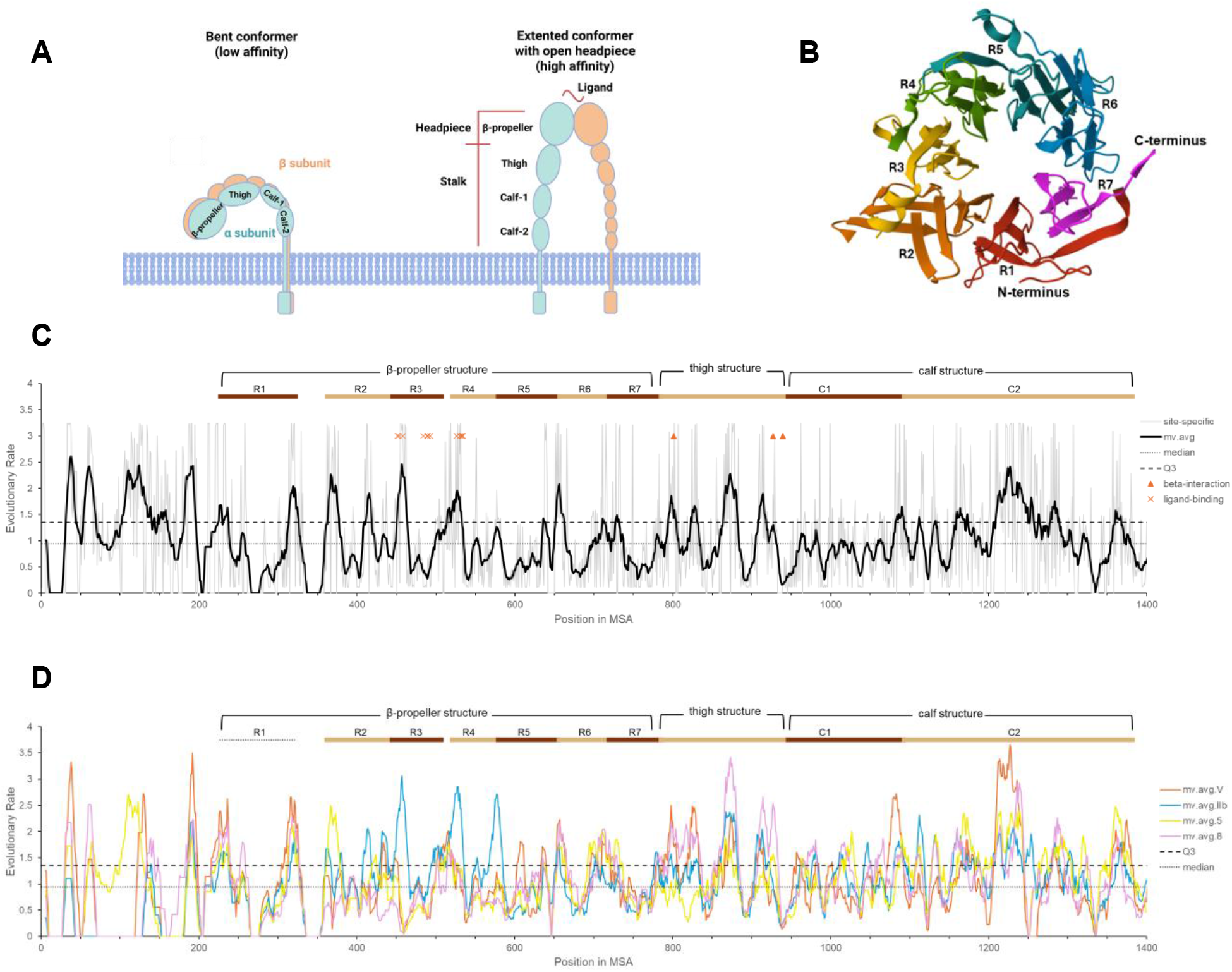
Structural organization and evolutionary rate variation of α-integrin ectodomains. **(A)** Schematic representation of integrin structure and conformational changes during activation transition. (**B**) Three-dimensional structure of the β-propeller headpiece composed of seven repeats. (**C**) Site-specific evolutionary rates estimated across all PS2 α-integrin sequences, with β-propeller repeats (R1–R7), thigh, and calf domains indicated. Moving averages (window size = 10 residues) are shown to highlight regional trends. Regions above the third quartile threshold are considered highly variable. (**D**) Paralog-specific evolutionary rate profiles for αIIb, α5, αV, and α8 based on orthologs from 28 vertebrate species.

We first estimated the evolutionary rates using all sequences across paralogs. As expected, regions of strong conservation are interspersed with more variable segments along the sequence, suggesting a mosaic pattern of structural constraint and functional flexibility. Using the third quartile (1.52 substitutions per million years) as a threshold, we identified highly variable regions (**Fig. 4C**). These regions are predominantly located within β-propeller repeats R2–R4, as well as in the thigh and calf-2 domains, suggesting that these structural elements are subject to relaxed constraint or lineage-specific functional diversification. Notably, several of these variable regions coincide with the RGD ligand-binding interface (limited to β-propeller domain) and the β-subunit interaction surfaces (involving β-propeller, thigh and calf-2 domains). These patterns indicate that functionally important sites may also be hotspots of adaptive divergence.

While evolutionary rates across all sequences reflect global conservation and divergence, paralog-specific rates capture lineage-specific evolutionary constraints across species. We therefore estimated evolutionary rates for each paralog separately using orthologs from the 28 vertebrate species. We observed pronounced paralog-specific variation across the α-integrin ectodomain (**Fig. 4D**). Although all paralogs share broadly similar patterns of conservation, several regions exhibit elevated evolutionary rates in one paralog relative to the others, indicating lineage-specific divergence. Notably, in the β-propeller region R3 – R5, αIIb exhibits prominent rate peaks, which are not mirrored in other paralogs. In contrast, α8 displays a distinct spike in the thigh region, and αV shows elevated rates in the calf segments. These localized differences suggest that, despite a shared structural framework, individual paralogs have undergone differential evolutionary pressures at specific sites.

### PS2 α-integrins evolved via neofunctionalization and specialization from a generalist ancestor

PS2 α-integrin paralogs pair with distinct subunits to form functional heterodimers and exhibit specialized ligand affinities and tissue-specific expression patterns. To pinpoint the evolutionary branches at which key functional shifts occurred, we reconstructed ancestral states using an MP approach based on the functional annotations of human paralogs (**Table 1**).

**Table 1.** Functional traits of extant and inferred ancestral α-integrin.

|  | Heterodimer forming |  |  | Ligand binding |  |  | Cells expressed |  |  |  |
| --- | --- | --- | --- | --- | --- | --- | --- | --- | --- | --- |
| | $\beta 1$ | $\beta 3$ | $\beta 5$ | Fn | Fb | Vn | PI | Im | Ms | Ep |
| human $\alpha IIb$ | 0 | 1 | 0 | 1 | 1 | 1 | 1 | 0 | 0 | 1 |
| human $\alpha 5$ | 1 | 0 | 0 | 1 | 0 | 0 | 1 | 1 | 1 | 1 |
| human $\alpha V$ | 1 | 1 | 1 | 1 | 1 | 1 | 1 | 1 | 1 | 1 |
| human $\alpha 8$ | 1 | 0 | 0 | 1 | 0 | 1 | 0 | 0 | 1 | 1 |
| MRCA | 1 * | 0 * | 0 | 1 | 0 * | 1 | 1 | 0 | 0 * | 1 |
| Anc_8_5_V | 1 | 0 | 0 | 1 | 0 | 1 | 1 | 0 | 1 | 1 |
| Anc_5_V | 1 | 0 | 0 | 1 | 0 | 1 | 1 | 1 | 1 | 1 |
| Anc_IIb | 0 | 1 | 0 | 1 | 1 | 1 | 1 | 0 | 0 | 1 |
| Anc_8 | 1 | 0 | 0 | 1 | 0 | 1 | 0 | 0 | 1 | 1 |
| Anc_5 | 1 | 0 | 0 | 1 | 0 | 0 | 1 | 1 | 1 | 1 |
| Anc_V | 1 | 1 | 1 | 1 | 1 | 1 | 1 | 1 | 1 | 1 |

Fn (fibronectin), Fb (fibrinogen), Vn (vitronectin)
PI (platelets/thrombocytes), Im (immune system cells), Ms (muscle tissue), Ep (epithelia tissue)
\* indicate ancestral states resolved based on additional α-integrin data in Hughes's 2001 paper

Our reconstruction indicates that the MRCA of vertebrate PS2 α-integrins originally paired with only β1. This ancestral state was retained in α5, αV, and α8 but lost in αIIb. Over evolutionary time, additional β-subunits became available as potential pairing partners for newly diverged α-subunits. Accordingly, the αIIb lineage evolved specific pairing with β3, while αV independently expanded its repertoire to interact with both β3 and β5, reflecting a significant gain in heterodimerization versatility.

The MRCA could bind to Fn and Vn in ECM. While this capacity was largely conserved, the α5 lineage lost Vn binding to become Fn-specific. Notably, the MRCA lacked Fb affinity, suggesting that Fb binding in αIIb and αV is acquired through independent, convergent evolution.

The expression profile of the MRCA was characterized by presence in platelets/thrombocytes and epithelial tissues, an ancestral state retained by most descendants except for α8. In contrast, expressions in immune cells and muscle tissue were absent in the MRCA. Muscle tissue expression was acquired early in the common ancestor of the α8, α5, and αV lineages (Anc_8_5_V), whereas immune cell expression emerged later in the common ancestor of the α5 and αV lineages (Anc_5_V).

These results reveal an evolutionary trajectory of neofunctionalization and specialization within the PS2 α-integrin subfamily. The MRCA was likely a broadly expressed receptor in epithelia and platelets/thrombocytes with versatile ECM-binding capacity (Fn and Vn) and forming dimers with the β1 subunit. Following early duplication events that gave rise to Anc_8_5_V and Anc_5_V, this lineage expanded its expression to additional tissues, including muscle and immune cells. Subsequent divergence of Anc_5_V into α5 and αV resulted in distinct functional outcomes. The α5 lineage largely retained ancestral features but lost Vn binding, becoming a specialized, high-affinity receptor for fibronectin required for matrix assembly. In contrast, αV evolved into the most functionally versatile paralog, acquiring the ability to pair with β3 and β5 and independently gaining Fb binding, consistent with its broad roles in angiogenesis, tumor progression, and TGF-β activation. The α8 lineage reflects a transition toward tissue-restricted function; although it retained ancestral β1 pairing and Fn/Vn binding, it lost platelet/thrombocyte expression and became primarily associated with muscle and epithelial tissues, supporting roles in development and fibroblast differentiation. Finally, the αIIb lineage underwent the most pronounced specialization, losing β1 pairing in favor of a dedicated β3 interaction and independently acquiring Fb binding. These changes underpin its platelet/thrombocyte-restricted role in Fb-dependent clot formation, a major functional transition associated with vertebrate hemostasis.

### Integrative analysis reveals genetic bases underlying functional specialization

Although all four PS2 α-integrin paralogs recognize the conserved RGD motif, their ligand specificities differ. In the previous section, our results indicate that Fn and Vn binding are ancestral traits retained across paralogs, whereas Fb binding represents a neofunction independently acquired by the αIIb and αV lineages (**Table 1**). To identify genetic basis of this convergent evolution, we mapped sequence variation to functional traits across reconstructed ancestral sequences, enabling identification of residues associated with lineage-specific functional divergence. Mapping to ancestral nodes, rather than relying solely on extant human sequences, allows discrimination between conserved ancestral features and derived adaptations, while minimizing confounding effects of species-specific mutations.

We focused on a segment corresponding to human αIIb residues 255–263 (NP_000410), αV residues 242–249 (NP_002201), α5 residues 262–269 (NP_002196), and α8 residues 257–264 (NP_003629), where αIIb exhibits accelerated evolutionary rates (**Fig. 4C**). This segment forms part of a ligand-binding specificity–determining loop, in which residues create a “cap” that shapes ligand interaction together with the paired β-subunit (**Fig. 5A-B**) [21, 22]. Because αIIbβ3 and αVβ3 are the only integrin heterodimers capable of binding fibrinogen (Fb), and our earlier analyses indicate that this function arose independently in αIIb and αV, we investigated the sequence changes associated with this convergent neofunctionalization.

**Figure 5.**
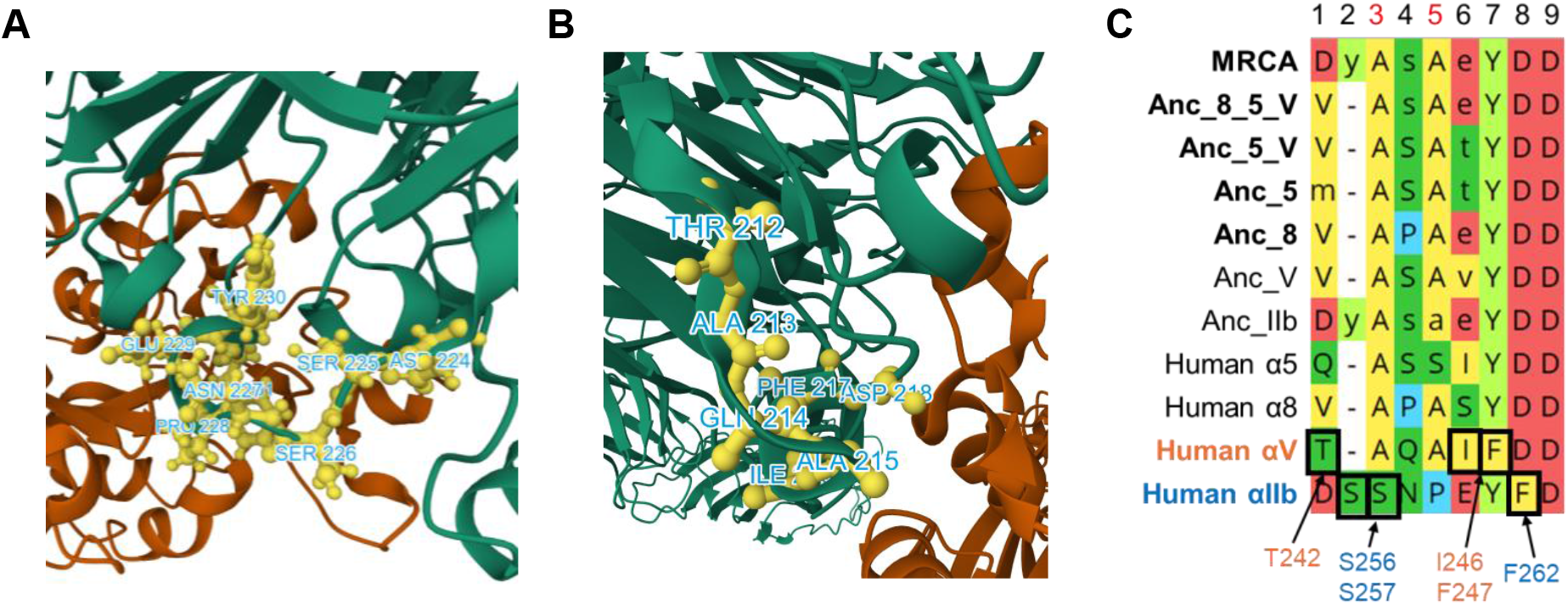
Structural and sequence basis of ligand-binding divergence. **(A, B)** Structural localization of the ligand-binding loop in human αIIbβ3 (A) and αVβ3 (B) integrins. Residues within the analyzed segment are shown in ball-and-stick representation and labeled; the α-subunit is colored orange, and the β-subunit is colored green. (**C**) Alignment of ancestral and extant sequences highlighting lineage-specific changes across the analyzed segments. Residues likely contributing to Fb binding acquisition are annotated.

Specifically, we compared residues in αIIb and αV with those in the MRCA, Anc_8_5_V, Anc_5, and Anc_8 sequences, which we previously inferred to lack Fb binding (**Table 1**). In αIIb, positions 256 and 257 reflect a lineage-specific S insertion and A→S substitution, respectively (**Fig. 5C**). These changes convert a nonpolar hydrophobic region in the ancestral sequences into two polar, hydrophilic residues. In addition, at position 262, a D→F substitution replaces a short acidic side chain with a bulky aromatic residue, potentially altering the physicochemical properties of the binding interface. In contrast, αV retains ancestral residues at these corresponding positions. Instead, substitutions associated with functional divergence in αV occur at distinct sites, including D→T at position 242, E→I at position 246, and Y→F at position 247. These results indicate that αIIb and αV achieved similar functional outcomes through different structural modifications, consistent with independent evolutionary trajectories and convergent neofunctionalization.

## Discussion

Integrins are ancient cell-surface receptors that have diversified to mediate complex cell–ECM interactions. In this study, we reconstruct the evolutionary trajectory of vertebrate PS2 α-integrins, linking sequence evolution to functional divergence and therapeutic potential.

Our analysis incorporates a balanced representation of species spanning cartilaginous fishes to primates. This expanded taxonomic coverage addresses limitations of previous studies with uneven sampling and provides a more resolved view of PS2 α-integrin evolution. We date the MRCA of the four vertebrate PS2 α-integrin paralogs, αIIb, α5, α8, and αV, to approximately 762 Mya, within the Neoproterozoic era, a period associated with early metazoan diversification [23]. Diversification of these paralogs was completed prior to vertebrate radiation, with subsequent lineage-specific refinements shaping their modern functions. Among these paralogs, αIIb was the earliest to establish its lineage-specific identity, whereas αV was the most recently diverged, with these events separated by approximately 70 million years, reflecting a prolonged process of diversification.

Ancestral reconstruction reveals that the PS2 MRCA was a β1-pairing receptor that bound Fn and Vn, with expression in platelets/thrombocytes and epithelia. Subsequent evolution drove functional partitioning. The αIIb lineage lost β1 to specialize in β3-mediated hemostasis, independently acquiring Fb binding. In contrast, αV expanded its repertoire, gaining β3 and β5 pairing flexibility and broad ligand affinity. Meanwhile, α5 specialized in Fn-binding, and α8 evolved toward tissue-restricted roles. This transition from a generalist ancestral state to lineage-specific specialization exemplifies neofunctionalization following gene duplication.

We observed distinct, domain-partitioned evolutionary strategies between the two Fb-binding paralogs, αIIb and αV. αIIb exhibits accelerated evolution in its ligand-binding β-propeller regions, reflecting adaptive optimization across species for rapid, shear-dependent fibrinogen recognition essential for clotting. Conversely, αV shows elevated evolutionary rates in its calf domains, favoring heterodimerization flexibility with multiple β subunits. Thus, αIIb evolved as a specialist through ligand-interface optimization, whereas αV evolved as a generalist through structural expansion. By mapping these evolutionary substitutions onto extant and ancestral sequences, we identified lineage-specific sites at the Fb-binding interface that likely facilitated the independent acquisition of this neofunction.

These findings have significant translational implications. The functional specialization and structural constraints of αIIb facilitate selective targeting, whereas the versatility and redundancy of αV complicate it. The lineage-specific sites identified at the Fb-binding interface serve as high-priority candidates for the development of precision therapeutics. This evolution-informed framework not only elucidates the diversification of the PS2 integrin subfamily but also provides a rational basis for designing selective antagonists that minimize off-target effects, a strategy broadly applicable to other complex and highly conserved protein families.

Despite the insights provided by this study, several limitations must be considered. First, our analysis focused exclusively on the PS2 α-integrin subfamily. While this allowed for a high-resolution view of RGD-binding evolution, including the broader α-integrin subunits would be necessary to reconstruct the full evolutionary landscape of the integrin superfamily and identify potential cross-clade functional redundancies. Furthermore, while we focused on the α-subunits as the primary drivers of ligand specificity, incorporating the pairing β-subunits into a co-evolutionary analysis would likely reveal deeper insights into the structural constraints of heterodimerization and how compensatory mutations across the interface maintain functional integrity. Lastly, our taxonomic sampling does not include lamprey. Although genomic data are available, the divergence of α-integrin-like proteins in this lineage is such that they cannot be accurately classified into precise vertebrate subtypes using current phylogenetic models. Future studies leveraging improved assembly and annotation of early vertebrate genomes may help resolve these basal nodes and further refine the timing of the PS2 diversification.

## Notes

### Competing Interest Statement

The authors have declared no competing interest.

### Funding

- National Institutes of Health: 1R01LM013438

