## Supplementary Table 1 for "From Generalist to Specialist: Evolution of PS2 α-integrins and Implications for Drug Targeting"

**Supplementary Table 1.** Species and sequences used in the analyses

| | | Common name | Taxa name | Accession # | Start | End |
|---|---|---|---|---|---|---|
| αV | | | | | | |
| | 1 | Bottlenose dolphin | Tursiops truncatus | XP_004317322 | 31 | 991 |
| | 2 | Gray whale | Eschrichtius robustus | XP_068400917 | 31 | 990 |
| | 3 | Elk | Cervus canadensis | XP_043344172 | 31 | 991 |
| | 4 | Red deer | Cervus elaphus | XP_043750716 | 31 | 991 |
| | 5 | Horse | Equus przewalskii | XP_008528799 | 31 | 990 |
| | 6 | Zebra | Equus quagga | XP_046513782 | 31 | 990 |
| | 7 | Dog | Canis lupus familiaris | XP_038302681 | 25 | 984 |
| | 8 | Wolf | Canis lupus baileyi | XP_072666955 | 35 | 994 |
| | 9 | Fruit bat | Artibeus jamaicensis | XP_036992717 | 31 | 991 |
| | 10 | Rabbit | Oryctolagus cuniculus | XP_002712375 | 30 | 990 |
| | 11 | European hare | Lepus europaeus | XP_062044852 | 30 | 990 |
| | 12 | European shrew | Sorex araneus | XP_054976855 | 44 | 1004 |
| | 13 | Rat | Arvicanthis niloticus | XP_034350141 | 31 | 986 |
| | 14 | Mouse | Mus musculus | NP_032428 | 31 | 987 |
| | 15 | Human | Homo sapiens | NP_002201 | 31 | 991 |
| | 16 | Chimpanze | Pan troglodytes | XP_515967 | 31 | 991 |
| | 17 | Koala | Phascolarctos cinereus | XP_020833200 | 47 | 1006 |
| | 18 | Common wombat | Vombatus ursinus | XP_027721735 | 47 | 1006 |
| | 19 | Bird | Phalacrocorax carbo | XP_064308112 | 91 | 1049 |
| | 20 | Tiger rattlesnake | Crotalus tigris | XP_039205902 | 20 | 980 |
| | 21 | Yellowpond turtle | Mauremys mutica | XP_044888647 | 22 | 981 |
| | 22 | European pond turtle | Emys orbicularis | XP_065269053 | 22 | 981 |
| | 23 | Fence lizard | Sceloporus undulatus | XP_042321005 | 23 | 984 |
| | 24 | Mimic poison frog | Ranitomeya imitator | XP_069604293 | 39 | 993 |
| | 25 | Zimmermann frog | Ranitomeya variabilis | XP_077140029 | 39 | 993 |
| | 26 | Zebrafish | Danio rerio | NP_001028893 | 25 | 987 |
| | 27 | Platypus | Ornithorhynchus anatinus | XP_028927467 | 18 | 977 |
| | 28 | Sawshark | Pristiophorus japonicus | XP_070731931 | 26 | 979 |
| α5 | | | | | | |
| | 1 | Bottlenose dolphin | Tursiops truncatus | XP_033722430 | 92 | 1049 |
| | 2 | Gray whale | Eschrichtius robustus | XP_068415706 | 92 | 1049 |
| | 3 | Elk | Cervus canadensis | XP_043302333 | 95 | 1051 |
| | 4 | Red deer | Cervus elaphus | XP_043737765 | 95 | 1051 |
| | 5 | Horse | Equus przewalskii | XP_070477652 | 96 | 1052 |
| | 6 | Zebra | Equus quagga | XP_046518283 | 96 | 1052 |
| | 7 | Dog | Canis lupus familiaris | XP_038293862 | 44 | 1000 |
| | 8 | Wolf | Canis lupus baileyi | XP_072653713 | 44 | 1000 |
| | 9 | Fruit bat | Artibeus jamaicensis | XP_037015552 | 94 | 1050 |
| | 10 | Rabbit | Oryctolagus cuniculus | XP_002711083 | 95 | 1050 |
| | 11 | European hare | Lepus europaeus | XP_062059520 | 97 | 1052 |
| | 12 | European shrew | Sorex araneus | XP_054981801 | 96 | 1052 |
| | 13 | Rat | Arvicanthis niloticus | XP_034372497 | 94 | 1048 |
| | 14 | Mouse | Mus musculus | NP_034707 | 45 | 1000 |
| | 15 | Human | Homo sapiens | NP_002196 | 42 | 996 |

| | | | | | |
|---|---|---|---|---|---|
| | 16 Chimpanze | Pan troglodytes | XP_016778761 | 91 | 1045 |
| | 17 Koala | Phascolarctos cinereus | XP_078001736 | 103 | 1077 |
| | 18 Common wombat | Vombatus ursinus | XP_027713858 | 101 | 1053 |
| | 19 Bird | Phalacrocorax carbo | XP_064330440 | 34 | 999 |
| | 20 Tiger rattlesnake | Crotalus tigris | XP_039200678 | 39 | 1000 |
| | 21 Yellowpond turtle | Mauremys mutica | XP_044850702 | 21 | 985 |
| | 22 European pond turtle | Emys orbicularis | XP_065275268 | 25 | 989 |
| | 23 Fence lizard | Sceloporus undulatus | XP_042305701 | 29 | 981 |
| | 24 Mimic poison frog | Ranitomeya imitator | XP_069614813 | 25 | 987 |
| | 25 Zimmermann frog | Ranitomeya variabilis | XP_077154036 | 21 | 983 |
| | 26 Zebrafish | Danio rerio | NP_001004288 | 33 | 1001 |
| | 27 Platypus | Ornithorhynchus anatinus | XP_028929918 | 22 | 982 |
| | 28 Sawshark | Pristiophorus japonicus | XP_070725425 | 78 | 1036 |
| α8 | | | | | |
| | 1 Bottlenose dolphin | Tursiops truncatus | XP_033708369 | 38 | 1008 |
| | 2 Gray whale | Eschrichtius robustus | XP_068418064 | 38 | 1008 |
| | 3 Elk | Cervus canadensis | XP_043334344 | 38 | 1008 |
| | 4 Red deer | Cervus elaphus | XP_043738642 | 38 | 1008 |
| | 5 Horse | Equus przewalskii | XP_070457181 | 38 | 1008 |
| | 6 Zebra | Equus quagga | XP_046535364 | 38 | 1008 |
| | 7 Dog | Canis lupus familiaris | XP_038514274 | 38 | 1008 |
| | 8 Wolf | Canis lupus baileyi | XP_072681792 | 38 | 1008 |
| | 9 Fruit bat | Artibeus jamaicensis | XP_036997190 | 38 | 1008 |
| | 10 Rabbit | Oryctolagus cuniculus | XP_002717473 | 38 | 1008 |
| | 11 European hare | Lepus europaeus | XP_062066487 | 38 | 1008 |
| | 12 European shrew | Sorex araneus | XP_012787480 | 27 | 997 |
| | 13 Rat | Arvicanthis niloticus | XP_034366862 | 38 | 1008 |
| | 14 Mouse | Mus musculus | NP_001001309 | 38 | 1008 |
| | 15 Human | Homo sapiens | NP_003629 | 39 | 1009 |
| | 16 Chimpanze | Pan troglodytes | XP_507671 | 39 | 1009 |
| | 17 Koala | Phascolarctos cinereus | XP_020843163 | 39 | 1009 |
| | 18 Common wombat | Vombatus ursinus | XP_027703749 | 39 | 1009 |
| | 19 Bird | Phalacrocorax carbo | XP_064300768 | 36 | 1002 |
| | 20 Tiger rattlesnake | Crotalus tigris | XP_039187554 | 42 | 1007 |
| | 21 Yellowpond turtle | Mauremys mutica | XP_044860657 | 84 | 1049 |
| | 22 European pond turtle | Emys orbicularis | XP_065254887 | 87 | 1052 |
| | 23 Fence lizard | Sceloporus undulatus | XP_042332726 | 28 | 993 |
| | 24 Mimic poison frog | Ranitomeya imitator | XP_069585550 | 26 | 998 |
| | 25 Zimmermann frog | Ranitomeya variabilis | XP_077124359 | 26 | 998 |
| | 26 Zebrafish | Danio rerio | XP_021322596 | 37 | 1004 |
| | 27 Platypus | Ornithorhynchus anatinus | XP_028933634 | 24 | 995 |
| | 28 Sawshark | Pristiophorus japonicus | XP_070737431 | 23 | 989 |
| αIIb | | | | | |
| | 1 Bottlenose dolphin | Tursiops truncatus | XP_073654545 | 32 | 994 |
| | 2 Gray whale | Eschrichtius robustus | XP_068386267 | 32 | 990 |
| | 3 Elk | Cervus canadensis | XP_043322763 | 32 | 988 |
| | 4 Red deer | Cervus elaphus | XP_043760773 | 32 | 988 |

| | | | | | |
|---|---|---|---|---|---|
| 5 | Horse | Equus przewalskii | XP_008518827 | 104 | 1093 |
| 6 | Zebra | Equus quagga | XP_046533559 | 104 | 1093 |
| 7 | Dog | Canis lupus familiaris | NP_001003163 | 32 | 993 |
| 8 | Wolf | Canis lupus baileyi | XP_072636954 | 81 | 1042 |
| 9 | Fruit bat | Artibeus jamaicensis | XP_037018088 | 33 | 987 |
| 10 | Rabbit | Oryctolagus cuniculus | NP_001075534 | 32 | 989 |
| 11 | European hare | Lepus europaeus | XP_062030824 | 32 | 988 |
| 12 | European shrew | Sorex araneus | XP_004621085 | 32 | 993 |
| 13 | Rat | Arvicanthis niloticus | XP_034360285 | 32 | 992 |
| 14 | Mouse | Mus musculus | NP_034705 | 32 | 989 |
| 15 | Human | Homo sapiens | NP_000410 | 32 | 994 |
| 16 | Chimpanze | Pan troglodytes | XP_024206013 | 101 | 1063 |
| 17 | Koala | Phascolarctos cinereus | XP_020853356 | 32 | 1009 |
| 18 | Common wombat | Vombatus ursinus | XP_027729022 | 32 | 1009 |
| 19 | Bird | Phalacrocorax carbo | XP_064329275 | 26 | 986 |
| 20 | Tiger rattlesnake | Crotalus tigris | XP_039187229 | 34 | 963 |
| 21 | Yellowpond turtle | Mauremys mutica | XP_044855696 | 35 | 1001 |
| 22 | European pond turtle | Emys orbicularis | XP_065278640 | 35 | 1001 |
| 23 | Fence lizard | Sceloporus undulatus | XP_042330649 | 31 | 993 |
| 24 | Mimic poison frog | Ranitomeya imitator | XP_069607515 | 16 | 983 |
| 25 | Zimmermann frog | Ranitomeya variabilis | XP_077108705 | 17 | 984 |
| 26 | Zebrafish | Danio rerio | NP_001003857 | 24 | 983 |
| 27 | Platypus | Ornithorhynchus anatinus | XP_028930901 | 33 | 997 |
| 28 | Sawshark | Pristiophorus japonicus | XP_070720604 | 33 | 1003 |
| outgroup | | | | | |
| 1 | Fruit fly | Drosophila melanogaster | P12080 | 32 | 1341 |
| 2 | Round worm | Caenorhabditis elegans | NP_498948 | 26 | 1152 |
